# Chronic cocaine exposure negatively impacts Long-COVID-like outcomes produced by the SARS-CoV-2 spike protein in the rat

**DOI:** 10.64898/2026.06.02.729575

**Authors:** Sarah E. Davis, Danielle R. Stern, Saadet Inan, Emily Vu, Daniel Lopez, Flourish Anwuri, Marco G. Ghilotti, Joseph J. Meissler, Ellen M. Unterwald

**Affiliations:** Center for Substance Abuse Research and Department of Neural Sciences, Lewis Katz School of Medicine at Temple University, Philadelphia, PA, USA

**Keywords:** Long-COVID, cocaine, SARS-CoV-2 spike protein, neuroinflammation, mechanical allodynia, impulsivity

## Abstract

Acute COVID-19 outcomes are exacerbated by substance use, however, the impact of substance use on Long-COVID is unknown. Here, we investigated the impact of chronic cocaine administration on spike-induced Long-COVID-like outcomes in the rat. Rats received intermittent chronic cocaine administration and a single intravenous injection of the SARS-CoV-2 spike protein. Two months following spike administration, Long-COVID-like outcomes were assessed. Exposure to spike protein in the presence of cocaine produced a persistent reduction in weight gain as compared with controls or spike protein alone. Further, cocaine-treated rats exposed to spike had lower withdrawal thresholds compared to control animals as well as their own baseline, suggesting increased pain sensitivity. Spike and/or cocaine increased the ratio of interleukin-6 (IL-6) to interleukin-10 (IL-10) levels in the hippocampus, indicating a shift towards a proinflammatory state. Paw withdrawal thresholds were positively correlated with IL-10 levels in the hippocampus and prefrontal cortex. Regarding olfaction, rats exposed to spike spent less time sniffing an odor attractant. Cocaine produced an anxiolytic-like phenotype during the elevated plus maze test. Further analysis of behaviors on the maze revealed that the latency to enter the open arms was shorter in rats exposed to spike or cocaine, suggesting a possible impulsive-like phenotype in these animals. These findings demonstrate the negative impact of cocaine on Long-COVID-like outcomes suggesting a need for increased clinical observations of people with co-occurring Long-COVID and cocaine use disorder.

**Graphical Abstract:** 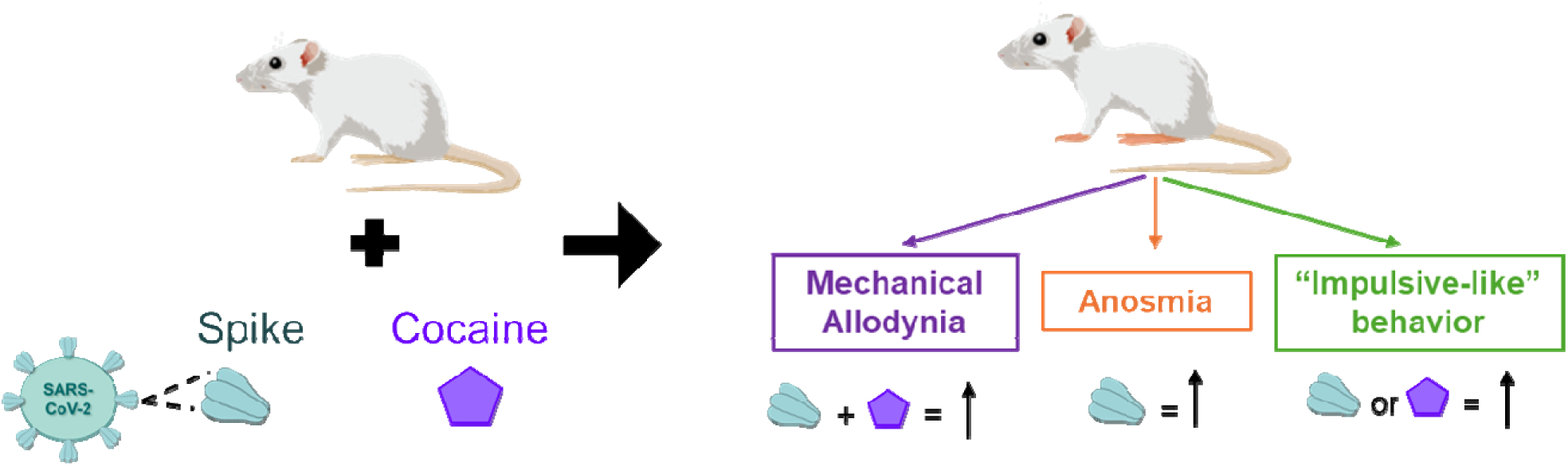

## Introduction

Estimates predict that 5-6% of adults who contracted COVID-19 during 2020 – 2024 developed post contemporary corona virus disease 2019 condition, also known as Long-COVID (Mandel et al. 2025). Symptoms of Long-COVID prominently feature neurologic manifestations including anxiety, depression, and cognitive disturbances (“brain fog”). Importantly, substance use has been linked to a higher risk for developing COVID-19 disease, as well as more severe adverse outcomes (morbidity/mortality) associated with acute COVID-19 disease (Hossain et al. 2020; Allen et al. 2021; Wang et al. 2022; Ramakrishnan et al. 2022). While 82% of Americans have received at least one COVID-19 vaccine dose, individuals with Long-COVID and a co-occurring substance use disorder have low vaccination rates; only 4.7% of 157,967 people with a co-occurring substance use disorder and Long-COVID were vaccinated against COVID-19 (World Health Organization 2023; Tsai et al. 2024). Despite the drastically low vaccination rate in this population, the impact of co-occurring cocaine-use on Long-COVID outcomes has not been investigated.

Preclinical reports have identified a potential role for the SARS-CoV-2 spike protein in mediating the neurological deficits associated with Long-COVID. The wildtype spike protein S1 subunit persists in mouse brain for at least 28 days post injection (Rong et al. 2024). Mice that received intracerebroventricular infusion of the spike protein express latent anxiety-like behaviors and deficits in object memory 30 and 45 days post spike exposure (Fontes-Dantas et al. 2023; Rong et al. 2024). These behavioral deficits coincide with elevated expression of neuroinflammatory markers reflecting similar clinical observations noted in persons with Long-COVID (Bansal et al. 2024). Cocaine and spike independently alter inflammatory responses. For example, serum levels of the anti-inflammatory cytokine IL-10 are lower in people who use cocaine (Fox et al. 2012; Moreira et al. 2016). Similarly, serum levels of IL-10 are reduced in people with Long-COVID (Carlini et al. 2023). Therefore, we hypothesized that chronic cocaine administration would worsen spike-induced Long-COVID outcomes and that this may be mediated by neuroinflammatory cytokines such as IL-10.

To test the hypothesis that cocaine negatively impacts Long-COVID outcomes, male Sprague Dawley rats were chronically exposed to cocaine, injected intravenously (iv) with SARS-CoV-2 wildtype spike protein, and then maintained on intermittent cocaine for 2 additional months, after which anxiety-like behaviors and memory were tested. In addition, mechanical allodynia was assessed, as musculoskeletal pain has been reported by persons with Long-COVID (Khoja et al. 2024). Anosmia is a frequent symptom of acute COVID-19 and Long-COVID (Kim et al. 2024). A retrospective study found a higher risk for anosmia in persons with Long-COVID and a co-occurring substance use disorder compared to those without a substance use disorder (Tsai et al. 2024). Therefore, olfaction was assessed by recording time spent sniffing a neutral or odor attractant (Yang and Crawley 2009). Finally, neuroinflammatory proteins and expression of genes associated with the aforementioned behaviors were measured. This study presents the first insights into the impact of co-occurring chronic cocaine and SARS-CoV-2 spike protein on neurologic functions and inflammation in a preclinical rat model.

## Methods

### 1. Study Design – Animals and Injection Schedule

Thirty-eight adult male Sprague Dawley rats (Charles River Laboratories Inc.) were used in the present study. Rats were housed in pairs with ad libitum access to food and water on a 12-hour light/dark cycle. Body weights were recorded daily throughout the study. Protocols were approved by the Temple University Institutional Animal Care and Use Committee. After acclimation to the facility for one week, rats were arbitrarily assigned to one of four treatment groups: control (saline iv + saline ip, spike alone (spike iv + saline ip), cocaine alone (cocaine ip + saline iv), or combination spike and cocaine (cocaine ip + spike iv), where n = 9-10 rats/treatment group.

Rats received 3 injections of saline or cocaine (15 mg/kg ip) at one-hour intervals (ie, in a binge-pattern (Unterwald et al. 2001) daily for 7 days. The following day, rats were injected with saline or spike protein (500 ng/kg, iv) through the tail vein. Seventy-two hours after the saline or spike injection, rats entered a “3 days on/4 days off” injection schedule wherein they received saline or cocaine (15 mg/kg) three times daily for 3 days, followed by 4 days of no injections, until 45 days post spike injection. On days 46-54, rats received one injection/day of saline or cocaine (25 mg/kg) following the same 3 days on/4 days off protocol. Behavioral testing occurred on days 55 through 61. Rats were euthanized on days 62 – 63 post spike injection for tissue and serum collection.

### 2. Behavior Testing

Rats were assigned a random label for all behavior testing. All scoring was done by two independent scorers who were blinded to treatment groups assignment. Averages of the two scores were used for quantitative analysis. Rats were allowed to habituate in the testing room 30-45 minutes prior to behavioral testing. All tests were performed during the light phase.

#### 2.1 Mechanical Sensitivity by Von Frey Test

Paw withdrawal thresholds (PWT) were measured using Von Frey filaments to define mechanical sensitivity as previously described (Inan et al. 2018, 2024). PWTs were measured 3 days before the first cocaine or saline injection (Day −11) and again 1 day after the final cocaine or saline injection (Day 55). Rats were placed in square elevated plastic boxes with metal mesh floors and allowed to habituate to the box for 30 minutes prior to testing. A series of Von Frey filaments (Stoelting, Chicago, IL) with logarithmically incrementing stiffness were applied perpendicularly to the midplantar region of each hind paw. The monofilaments were applied with increasing force until the rat withdrew the paw. Each monofilament was applied three times, 10 seconds apart to a paw. The paw withdrawal threshold was defined as the lowest force that evoked a brisk withdrawal response to one of the three repetitive filament applications. If the rat withdrew the paw during one of the three trials, that force was recorded as the score for that paw. The right and left paws were tested 5 minutes apart. An average of the scores for the right and left paw was used for quantitative analysis.

#### 2.2 Novel Object Recognition and Open Field

On experimental day 58 or 60 post spike, rats were placed into an open field chamber and allowed to roam freely for 20 minutes while video recorded. The dimensions for the open field chamber were 45 cm x 45 cm x 43 cm. Light conditions for both the open field test and the novel object test were 60 lux at the center of the open field chamber. 3D-printed objects were used for the study (Davis 2026). Objects (familiar and novel) were cleaned with 70% ethanol before each test. The first 10 minutes of the 20-min habituation period were scored as the open field test. On the next day, two identical objects were placed in opposing corners of the chamber; rats were placed in the center and allowed to freely explore for 10 minutes. Rats were returned to their home cage for 30 minutes during which one object was switched for a ‘novel’ object. Rats were again allowed to explore and interact with the objects for 10 minutes. Total time spent with each object was measured. A discrimination index for the novel object was calculated as [(Time exploring novel – Time exploring familiar)/ (Time exploring novel + Time exploring familiar)] x 100. Video recordings for open field test were analyzed using EthoVision XT tracking software (Noldus, Leesburg, VA); time and entries in the center of the chamber (20 X 20 cm) and total distance traveled were measured to evaluate anxiety-like behaviors and locomotor activity, respectively.

#### 2.3 Elevated Plus Maze

Elevated plus maze testing was performed on day 61 or 62, as previously described by us (Denny et al. 2021). Light conditions were 150 lux open arms/135 lux closed arms. Rats were placed in the center of the maze and allowed to explore the elevated plus maze freely for 5 minutes while being video recorded from above.

#### 2.4 Y-maze

Y-maze testing was performed on day 56, as previously described (Mokkarala et al. 2025). Rats were placed in the center of the Y-maze (Maze Engineers, Skokie, IL) and allowed to freely explore for 5 minutes. Y-maze dimensions were 53 cm (end of arm to center), 10 cm (arm width), and 30 cm (wall height). Lighting conditions were 135 lux for the Y-maze arms. Spontaneous alternations were calculated and defined as three unique arms explored in succession. Moving triads were used to determine the total number of spontaneous alterations during the 5-min session. The percentage of spontaneous alternations was calculated as (# of spontaneous alternations/# of total entries – 2) x 100. Movement on the maze, including arm entries, was analyzed from video recordings using EthoVision XT tracking software (Noldus, Leesburg, VA) and validated by one independent observer who was blinded to treatment group.

#### 2.5 Olfaction

Olfaction testing was performed on day 58 or 59. The test was performed in clean chambers wherein a cotton swab previously moistened with either water or female rat urine was placed (Dinçkol et al. 2022). In the first phase of the olfaction test, rats were allowed to freely smell a cotton swab with water for 3 minutes, before being returned to their home cage for 45 minutes. Next, rats were placed in a new clean chamber with a cotton swab moistened with female rat urine and allowed to freely smell the cotton swab for 3 minutes. Different brand cotton swabs were used to facilitate visual differentiation between the two swabs during the two trials. Times spent sniffing the water and the female rat urine exposed cotton swabs were measured from video recordings. Rats that spent less than 10 seconds total sniffing (water + female rat urine) were excluded (N= 6/38 rats excluded).

### 3. Measurement of cytokines & chemokines in brain tissue and serum

Tissue and serum were collected on day 62 or 63 of the study. Fresh excised brain areas (olfactory bulb, prefrontal cortex, nucleus accumbens, hippocampus) were hemisected, frozen in 1.5 mL tubes and stored at −80 °C. Tissue samples for cytokine analysis were mechanically homogenized in 150 μL phosphate buffered saline (PBS) and 150 μL of Cell Lysis Buffer 2 (R&D Systems, Minneapolis, MN) and incubated for 30 minutes at room temperature. Samples were then centrifuged at 13,000 X g for 20 minutes, and supernatants were collected and stored at −80° C. Total protein concentrations (total protein mg/mL) were determined using a Pierce Bicinchoninic Acid (BCA) Protein Assay (Thermo Scientific, Rockford, IL). Whole blood samples for cytokine analysis rested for 1 hour at room temperature following collection to allow for clotting, then centrifuged at 2000 X g for 10 minutes at 4 °C. Serum was collected and stored at −80°C (Ehinger et al. 2021). Levels of rat IL-6 and IL-10 were determined from serum and tissue extracts using the Milliplex MAP Rat Cytokine/Chemokine Magnetic Bead Panel Luminex assay (Catalog # RECYTMAG-65k, Millipore, Burlington, MA). Samples were run in duplicate according to the kit instructions. The plate was read on a BioPlex 100 Luminex reader (BioRad Laboratories, Hercules, CA) using the BioPlex Manager 6.1 software. The pg/mL protein values used for quantitative analysis were determined by dividing the cytokine concentration (pg/mL extract) by their respective total protein mg/mL values.

### 4. RNA isolation and cDNA generation

Brain regions were dissected and immediately stored in 300 μL of RNA later solution (catalog # AM7024, Invitrogen) for 24 hours at 4°C then at −80°C until assayed for gene expression. RNA was isolated from the tissues using a Quick-RNA Miniprep kit (catalog # R1055, Zymo Research) after removal of the RNA later solution. Concentration and quality of RNA was determined using a Nanodrop 2000 spectrophotometer (Thermo Fisher). Samples were normalized to a concentration of 10 ng/μL RNA for complementary DNA (cDNA) synthesis. The cDNA synthesis was performed using a High-Capacity cDNA Reverse Transcription Kit (catalog #4374967, Applied Biosystems). Synthesis of cDNA was performed using the following temperature parameters: 25°C (10 minutes), 37°C (120 minutes), 85°C (5 minutes), 4°C (hold). Stable cDNA (concentration = 5 ng/μL) was stored at −20°C for later use.

### 5. Quantitative Polymerase Chain Reaction (qPCR)

Quantitative Real Time-PCR (qPCR) was performed using TaqMan Fast Advanced Master Mix (catalog # 4444557, Applied Biosystems), and TaqMan probes were used to quantify changes in gene expression (Thermo Fisher). TaqMan probe ID’s and corresponding cDNA (ng) used are listed in Table 1. Based on pilot studies, we determined *Hprt1* was an appropriate reference gene for the prefrontal cortex and *Gapdh* was appropriate for the remaining brain regions; *Hprt1* and *Gapdh* were not regulated by cocaine or spike in these regions. Experiments for the prefrontal cortex were tested in single plex while the remaining brain regions were tested in duplex. All target genes contained the FAM fluorophore and reference genes contained the VIC fluorophore. The qPCR reaction was carried out using a QuantStudio 3 Real-Time PCR system (Applied Biosystems) and a Comparative-CT-TaqMan relative quantification fast template for a 96-well plate (0.1 mL) using the Diomni^TM^ Design and Analysis 3.0 (RUO) software. Relative fold change in gene expression was calculated using the 2^-ΔΔCt^ method (Livak and Schmittgen 2001).

**Table 1.** Gene expression was measured using quantitative reverse transcription polymerase chain reaction (qRT-PCR). mRNA levels for targets in the hippocampus, nucleus accumbens, and olfactory bulb were normalized to *Gapdh*. mRNA of genes of interest in the prefrontal cortex were normalized to *Hprt1*. A cDNA concentration response analysis was performed for each individual target gene using qPCR, and based on cycle threshold values (Ct), an optimal amount of cDNA (ng) was chosen for each target. Primer IDs are listed for all targets and were purchased from Thermo Fisher Scientific.

| Target Gene | Protein Product | Full name | Primer ID | cDNA (ng) used | Brain Region(s) |
| --- | --- | --- | --- | --- | --- |
| *Gapdh* | GAPDH | Glyceraldehyde-3phosphate dehydrogenase | Rn01775763_g1 | - | Hippocampus, Nucleus Accumbens, Olfactory Bulb |
| *Hprt1* | HPRT-1 | Hypoxanthine-guanine phosphoribosyl transferase 1 | Rn1527840_m1 | - | Prefrontal Cortex |
| *Htr2a* | 5-HT<sub>2a</sub>R | Serotonin 2A receptor | Rn06357477_s1 | 5 | Prefrontal Cortex |
| *Htr2c* | 5-HT<sub>2c</sub>R | Serotonin 2c receptor | Rn06312433_m1 | 5 | Prefrontal Cortex |
| *c-fos* | c-Fos | Fos proto-oncogene | Rn02396759_m1 | 10 | Olfactory Bulb |
| *Gfap* | GFAP | Glial fibrillary acidic protein | Rn01253033_m1 | 5 | Hippocampus, Prefrontal Cortex |
| *Gria1* | GluA1 | Glutamate ionotropic receptor AMPA type subunit 1 | Rn06323759_m1 | 2.5 | Prefrontal Cortex, Nucleus Accumbens |
| *Grin2b* | GluN2B | NMDA receptor subtype 2B | Rn00680474_m1 | 5 | Prefrontal Cortex, Nucleus Accumbens |
| *Aif1* | Iba1 | Allograft inflammatory factor 1 | Rn03993468_g1 | 10 | Olfactory Bulb, Hippocampus, Prefrontal Cortex |

**Table 2.** Summary of results from qRT-PCR experiments that determined the effect of spike and/or cocaine on gene expression. Trends observed (not significant) are indicated by ns.

| Target (gene) | Protein Product | Brain Region | Two-way ANOVA main effects |
| --- | --- | --- | --- |
| *Htr2a* | 5-HT2aR | Prefrontal Cortex | cocaine ↑ (ns) |
| *Htr2c* | 5-HT2cR | Prefrontal Cortex | – |
| *Htr2a/Htr2c* | 5-HTaR/5-HT2cR | Prefrontal Cortex | – |
| *c-fos* | c-Fos | Olfactory Bulb | spike ↑ |
| *c-fos* | c-Fos | Prefrontal Cortex | spike ↑ |
| *Gfap* | GFAP | Prefrontal Cortex | – |
| *Gfap* | GFAP | Hippocampus | – |
| *Gria1* | GluA1 | Prefrontal Cortex | spike↑<br>cocaine ↑ |
| *Gria1* | GluA1 | Nucleus Accumbens | – |
| *Grin2b* | GluN2b | Prefrontal Cortex | cocaine ↑ |
| *Grin2b* | GluN2b | Nucleus Accumbens | cocaine ↓ |
| *Aif1* | Iba1 | Olfactory Bulb | – |
| *Aif1* | Iba1 | Prefrontal Cortex | cocaine ↑ (ns) |
| *Aif1* | Iba-1 | Hippocampus | spike ↓<br>cocaine ↓ (ns) |

### 6. Reagents

SARS-CoV-2 spike protein was purchased from Acro Biosystems (Newark, DE, catalog # SPN-C52H9). The spike trimer protein was derived from the wildtype (Wuhan) SARS-CoV-2 strain. Spike protein was reconstituted in ultra purified water for 1 hour and aliquoted for storage at −80°C. On the day of injection, an aliquot (10 μg/mL) was diluted in sterile saline to the working concentration (1 μg/uL), and 500 ng/kg body weight was injected iv in a volume of 0.5 mL/kg. Cocaine was generously provided by the National Institute on Drug Abuse (NIDA) Drug Supply Program. Cocaine was dissolved in sterile saline and injected ip in a volume of 1 ml/kg body weight.

### 7. Statistical Analysis

All data are presented as mean ± S.E.M. The statistical analyses were performed using GraphPad prism software 10. Threshold for significance was set as p < 0.05. Two-way analysis of variance (ANOVA) was used to determine the main effects of cocaine and spike when appropriate. Tukey’s post-hoc test was used when there were a significant main effect and significant interaction between treatment groups. Sidak’s post-hoc test was used to compare within-treatment groups when there was a significant main effect but no significant interaction between treatments (Wei et al. 2012). Statistically significant outliers were removed where appropriate (ROUT method, Q=1%).

## Results

### Cocaine and spike protein reduce body weight gain

Rats were weighed daily (starting weight range: 198-234 g). Mean weight gain (g) starting on day −10 for the four experimental groups are shown in Figure 1B. Two-way ANOVA with repeated measures revealed significant effects of time (F(1.864, 63.36) = 2309, p <0.0001), treatment (F(3,34) = 6.477, p = 0.0014), and interaction between time x treatment (F(5.591, 63.36) = 11.52, p < 0.0001). Post-hoc analysis using Dunnett’s multiple comparisons test shows significantly less weight gain between saline-cocaine vs saline-saline groups on days 32, 39, and 53 post spike injection, and significantly less weight gain between spike-cocaine vs saline-saline groups starting day 11 post spike injection which persisted through the remainder of the study.

**Figure 1:**
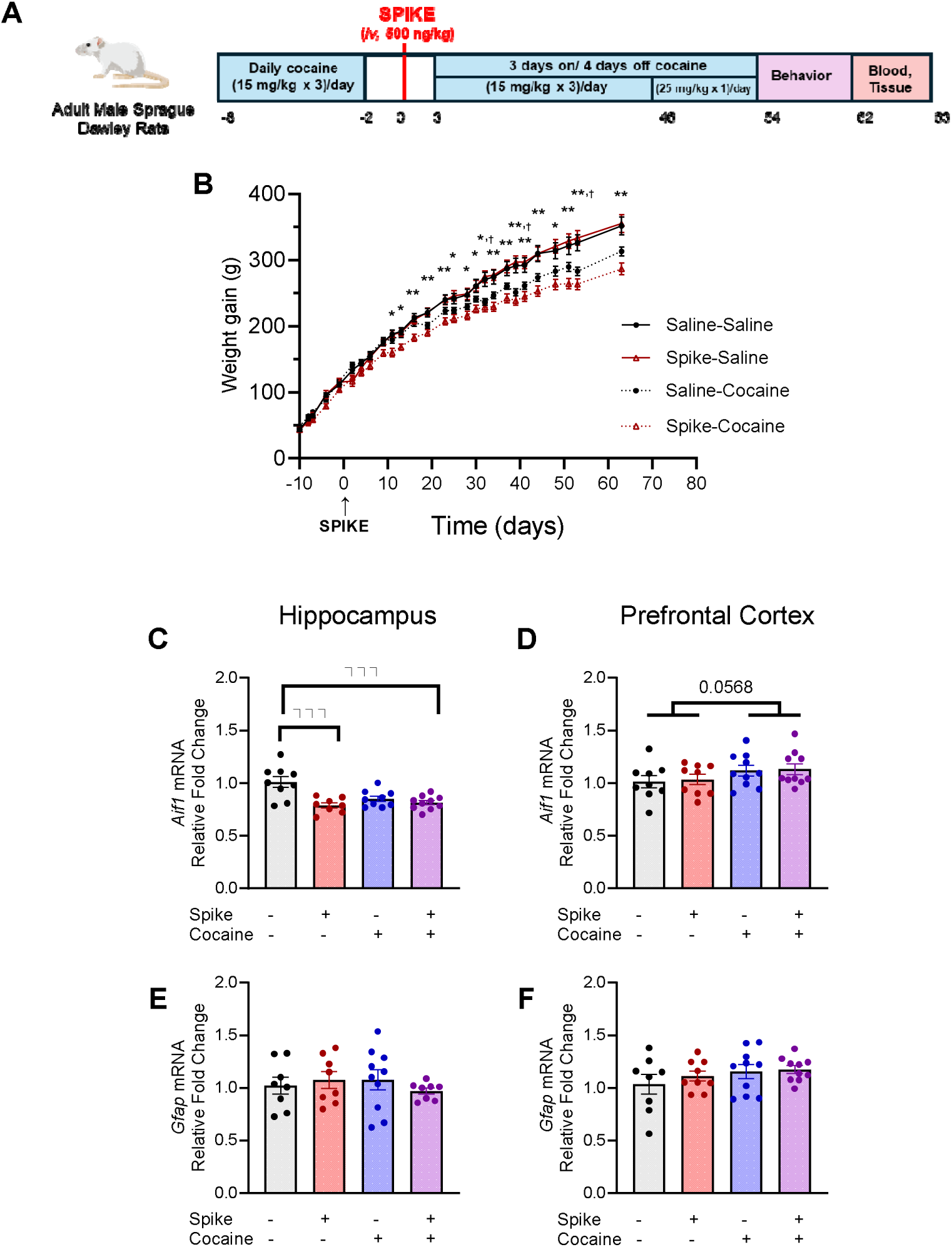
Spike and cocaine produced a persistent reduction in weight gain and altered *Aif1* expression in the brain. (A) Experimental timeline for the study. Rats received *i.p.* administration of cocaine or saline 3 times daily in a binge-pattern for 7 days. Rats then received *i.v.* injection of spike (500 ng/kg) or saline. Rats continued to receive cocaine or saline on a “3 days on - 4 days off” schedule through day 54. Behavioral experiments were performed between days 54 to 62. Rats were euthanized on days 62 and 63 and tissue and blood were collected for further study. (B) Cocaine reduced weight gain (g), and the reduction in weight gain was persistent and robust in the spike-cocaine group beginning 13 days after spike injection. Two-way ANOVA with repeated measures revealed main effects of time, treatment, and time x treatment. Significant results revealed by post-hoc analysis with Dunnett’s multiple comparisons test show reduced weight gain between saline-cocaine and saline-saline groups (^†^) as well as spike-cocaine and spike-saline groups (*). (C) Relative gene expression of *Aif1* was downregulated by spike and spike+cocaine in the hippocampus and (D) a trend for cocaine-induced upregulation of *Aif1* was observed in the prefrontal cortex. Two-way ANOVA revealed main effects of spike, spike x cocaine, and a trend for cocaine on *Aif1* expression the hippocampus. In the prefrontal cortex, two-way ANOVA revealed a trend for a main effect of cocaine on *Aif1* expression. (E) *Gfap* mRNA was unaltered by spike or cocaine in the hippocampus and (F) prefrontal cortex. Data are expressed as gene of interest relative to *Gapdh* or *Hprt1* mRNA in the hippocampus and prefrontal cortex respectively, and saline-saline rats are set as the control for calculated fold-changes. Data are expressed as means ± SEM. N = 8-10/group. Significance is denoted as p < 0.05, * or ^†^; p < 0.01, **; p < 0.001, ***.

### Spike and cocaine effects on microglial and astrocytic markers

To explore whether spike and/or cocaine altered microglial pathways, gene expression of *Aif1* (Iba1) and *Gfap* (GFAP) were quantified by qPCR in the hippocampus and prefrontal cortex. Regarding *Aif1* expression, two-way ANOVA revealed a significant interaction between spike x cocaine (F(1,32) = 7.763, p = 0.0089), as well as main effects of spike (F(1,32) = 15.40, p = 0.0004), and a trend for toward an effect for cocaine (F(1,32) = 4.098, p = 0.0513) in the hippocampus (Figure 1C). Further post-hoc analysis using Sidak’s multiple comparisons test showed that *Aif1* (Iba1) expression was downregulated in the spike-saline vs. saline-saline group (p = 0.0001) and spike+cocaine vs. saline-saline group (p = 0.0003). In the prefrontal cortex, there was a trend for cocaine to upregulate *Aif1* (two-way ANOVA: F(1,34) = 3.887, p = 0.0568) which did not reach statistical significance, with no significant effects of spike (F(1,34) = 0.1205, p = 0.7306) or interaction between spike x cocaine (F(1,34) = 0.0039, p = 0.9506) (Figure 1D). There were no significant effects of spike (F(1,30) = 0.027, p = 0.8708), cocaine (F(1,30) = 0.020, p = 0.8887) or interaction between spike x cocaine (F(1,30) = 1.873, p = 1.1813) on *Gfap* expression in the hippocampus (Figure 1E), or in the prefrontal cortex (spike (F(1,33) = 0.6205, p = 0.4365), cocaine (F(1,33) = 2.109, p = 0.1559), interaction (F(1,33) = 0.2108, p = 0.6492); Figure 1F). Together, these results suggest that spike or cocaine may regulate microgliosis in a region-specific manner.

### Spike and cocaine together produce mechanical allodynia

Musculoskeletal pain is a prevalent symptom of Long-COVID (Khoja et al. 2024). Here, the Von Frey test was used to assess potential changes in mechanical sensitivity. Paw withdrawal thresholds were measured prior to administration of cocaine or spike (baseline) and again 8 weeks following spike injection (Figure 2 A-B). Two-way ANOVA with repeated measures revealed a significant interaction between time x treatment (F(3,34) = 4.373, p = 0.0104). Post-hoc analysis using Tukey’s multiple comparison test revealed significantly lower paw withdrawal threshold between saline-saline vs. spike-cocaine animals at the 8-week time point (p = 0.0100), and between the baseline vs. 8-week time points for the spike-cocaine rats (p = 0.0226). There was also an increase between paw withdrawal thresholds of saline-saline rats at baseline vs. 8-weeks (p = 0.0113). We hypothesize this may be due to thicker padding on the rats’ paws due to increased body weight and aging which in turn may result in reduced mechanical sensitivity. These results indicate that co-exposure to spike and cocaine increased mechanical sensitivity 8 weeks after spike administration.

**Figure 2:**
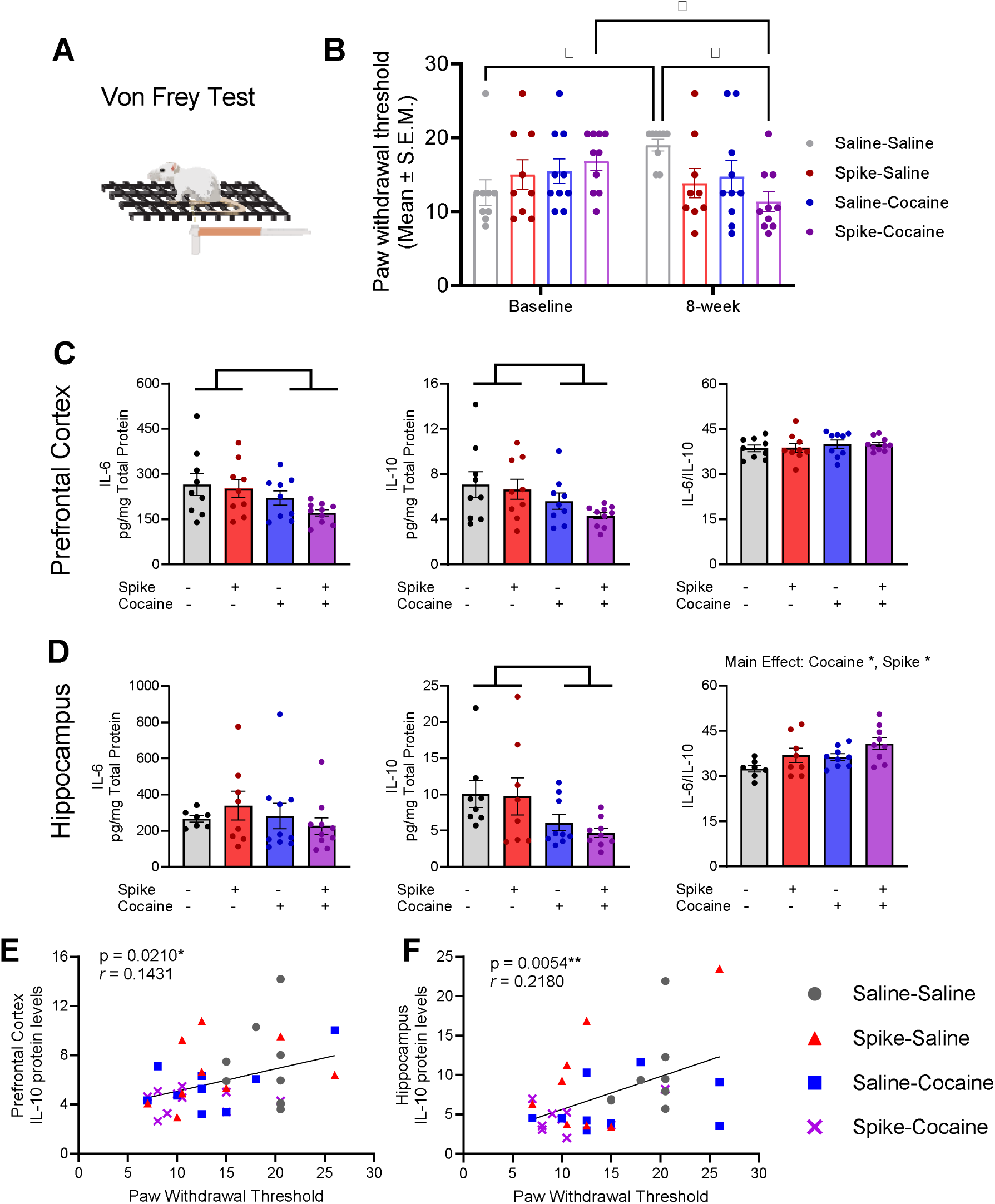
Rats exposed to spike protein plus cocaine showed enhanced mechanical allodynia which was associated with lower brain IL-10 levels. (A) Mechanical allodynia was assayed using Von Frey filaments at baseline (prior to receiving cocaine or spike) and 8 weeks after spike injection. (B) Spike-cocaine rats had lower paw withdrawal threshold scores (i.e., increased mechanical sensitivity) at 8 weeks relative to baseline thresholds and compared to saline-saline control rats at 8 weeks. Two-way ANOVA with repeated measures revealed a significant interaction between time x treatment. Significant results revealed by post-hoc analysis using Tukey’s multiple comparison test are shown (*). (C) Cocaine reduced IL-6 and IL-10 levels in the prefrontal cortex and (D) IL-10 in the hippocampus. Further, cocaine or spike increased the ratio of IL-6 to IL-10 in the hippocampus, indicating a shift toward a proinflammatory state. Two-way ANOVA revealed main effects of cocaine on IL-6 and IL-10 levels in the prefrontal cortex. In the hippocampus, two-way ANOVA revealed main effects of cocaine on IL-10 levels, and main effects of both spike and cocaine on the ratio of IL-6/IL-10 levels. € Paw withdrawal threshold scores significantly correlate with IL-10 levels in the prefrontal cortex (F) and hippocampus. Data (panels B-D) are expressed as means ± SEM. N = 8-10/group. Significance is denoted as p < 0.05, *; p < 0.01, **.

Elevation of the cytokines, interleukin-6 (IL-6) and IL-10, has been reported in the serum of persons with Long-COVID (Greene et al. 2024). Here, levels of cytokines were measured in the olfactory bulb, prefrontal cortex, hippocampus, and serum following long-term exposure to cocaine and spike. In the prefrontal cortex (Figure 2C), cocaine reduced IL-6 levels (two-way ANOVA revealed a significant effect of cocaine (F(1,33) = 5.608, p = 0.0239) but not spike (F(1,33) = 1.403, p = 0.2447) or interaction between spike x cocaine (F(1,33) = 0.4428, p = 0.5104). IL-10 levels were likewise lower in the prefrontal cortex of rats treated with cocaine (Fig 2C); two-way ANOVA revealed a significant effect of cocaine (F(1,33) = 5.584, p = 0.0242) but not spike (F(1,33) = 1.125, p = 0.2965), or interaction between spike x cocaine (F(1,33) = 0.3038, p = 0.5852). No difference in the ratio of IL-6 to IL-10 in the prefrontal cortex was found; two-way ANVOA showed no significant effects of cocaine (F(1,33) = 1.088, p = 0.3044), spike (F(1,33) = 0.0047, p = 0.9457), or interaction between spike x cocaine (F(1,33) = 0.017, p = 0.8970). In the hippocampus (Figure 2D), no significant differences in IL-6 levels were found (two-way ANOVA: cocaine (F(1,31) = 0.6073, p = 0.4417), spike (F(1,31) = 0.8815, p = 0.8815), or interaction (F(1,31) = 1.043, p = 0.3151)). Cocaine reduced IL-10 levels (F(1,30), p = 0.0096) in the hippocampus, although there was no significant effect of spike (F(1,30) = 0.2739, p = 0.6046) or interaction between spike x cocaine (F(1,30) = 0.1115, p = 0.7407) on IL-10. Analysis of the ratio of IL-6 to IL-10 in the hippocampus by two-way ANOVA revealed main effects of cocaine (F(1,29) = 6.456, p = 0.0167) and spike (F(1,29) = 5.039, p = 0.0326), but not interaction between spike x cocaine (F(1,29) = 0.0004, p = 0.9833) (Fig 2D). An increase in the ratio of IL-6 to IL-10 suggests a shift towards a proinflammatory state (You et al. 2011; Gholamnezhad et al. 2014; Moreira et al. 2016). IL-10 can reverse mechanical allodynia in rodent models of neuropathic pain (Laumet et al. 2020; Huang et al. 2022), hence we investigated the relationship between IL-10 and paw withdrawal thresholds in our model. There was a significant correlation between paw withdrawal threshold scores and IL-10 levels in both the prefrontal cortex (*r* = 0.1431, p = 0.0210)(Fig 2E) and the hippocampus (*r* = 0.2180, p = 0.0054) (Fig 2F). In serum, IL-6 and IL-10 were unaltered by treatment, however, the ratio of IL-6 to IL-10 levels were higher in cocaine-treated rats; two-way ANOVA revealed a significant effect of cocaine (F(1,32) = 4.406, p = 0.0438) (Figure S1).

We further explored changes in gene expression for proteins that have been associated with mechanical allodynia including GluA1 and GluN2B subunits of AMPA and NMDA glutamate receptors respectively (Su et al. 2015; Li et al. 2017; Fan et al. 2018; Wang et al. 2021). In the nucleus accumbens (Figure 3A), two-way ANOVA revealed no significant effect of spike (F(1,33) = 2.678, p = 0.1113), cocaine (F(1,33) = 0.8208, p = 0.3715), or interaction between spike x cocaine (F(1,33) = 0.2468, p = 0.6226) on *Grai1* (GluA1) expression. However, in the prefrontal cortex (Figure 3B), two-way ANOVA revealed significant effects of spike (F1,33) = 4.172, p = 0.0492) and cocaine (F(1,33) = 11.43, p = 0.0019), but no significant interaction between spike x cocaine (F(1,33) = 2.138, p = 0.1532) on *Gria1* gene expression. Sidak’s post-hoc test showed cocaine-exposed rats had higher *Gria1* relative to the saline-saline group (p = 0.0075), while the spike group had a trend for higher *Gria1* expression relative to the saline-saline group (p = 0.07). For *Grin2b* (GluN2B) expression in the nucleus accumbens (Figure 3C), two-way ANOVA revealed a significant interaction between spike x cocaine (F(1,31) = 6.443, p = 0.0164) and main effect of cocaine (F(1,31) = 6.005, p = 0.0201) but not spike (F(1,31) = 2.964, p = 0.0951). Sidak’s post-hoc test showed cocaine (p = 0.0045) and spike+cocaine (p = 0.0163) had lower *Grin2b* relative to the saline-saline group (Figure 3C). In the prefrontal cortex (Figure 3D), *Grin2b* expression was upregulated by cocaine; two-way ANOVA revealed significant main effect of cocaine (F(1,34) = 8.431, p = 0.0064), while spike (F(1,34) = 0.2896, p = 0.5940) and interaction between spike x cocaine (F(1,34) = 2.957, p = 0.0946) were not significant. Sidak’s post-hoc test showed the cocaine group had higher *Grin2b* relative to the saline-saline group (p = 0.0049; Figure 3D).

**Figure 3:**
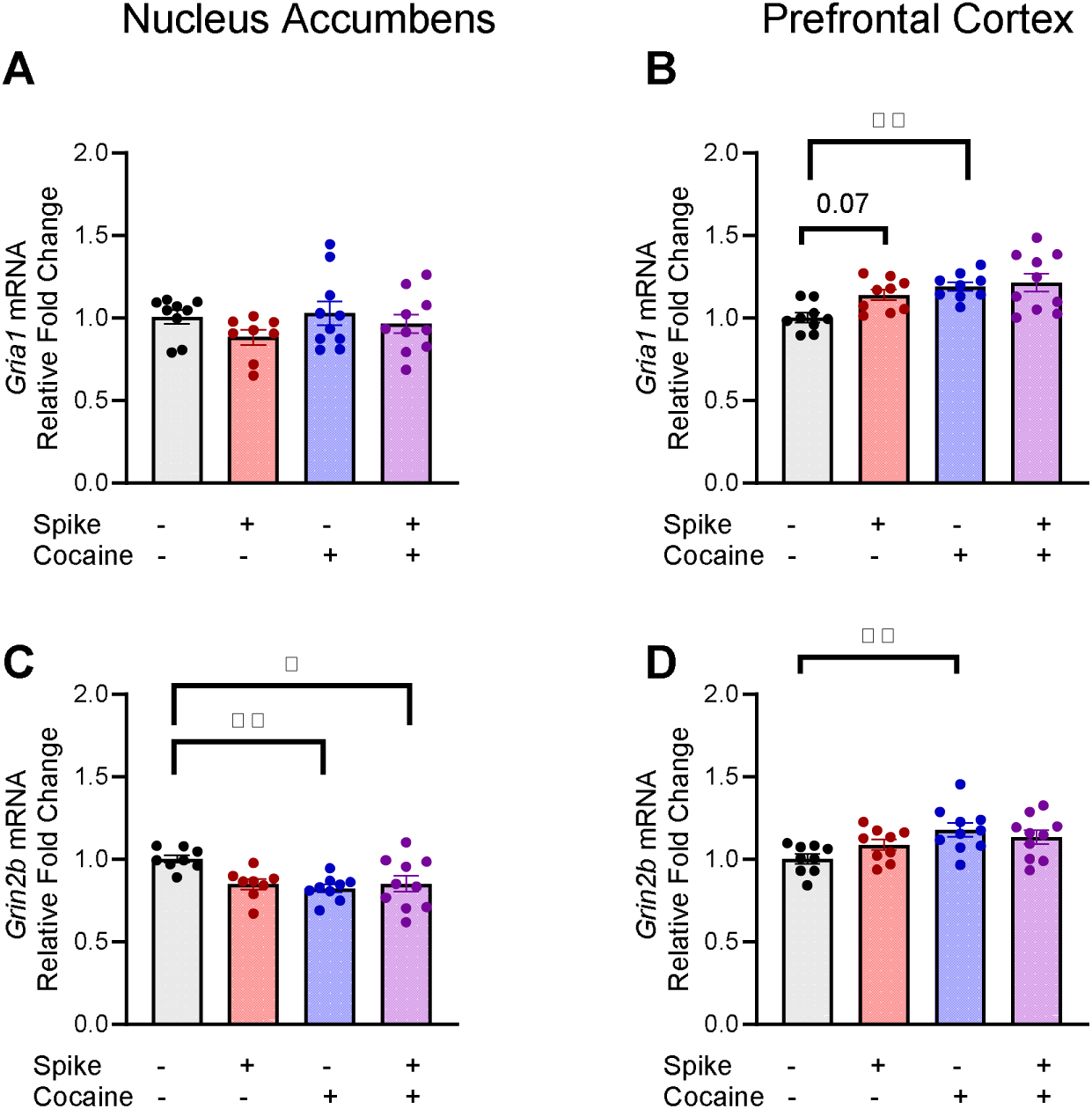
Gene expression of AMPA (*Gria1)* and NMDA (*Grib2)* receptor subunits were measured in the nucleus accumbens and prefrontal cortex of rats exposed to spike and/or cocaine. (A) Neither cocaine nor spike altered *Gria1* mRNA in the nucleus accumbens. (B) Spike or cocaine upregulated *Gria1* mRNA in the prefrontal cortex. Two-way ANOVA revealed main effects of cocaine and spike on *Gria1* expression in the prefrontal cortex. (C) *Grin2b* was downregulated by cocaine or spike+cocaine in the nucleus accumbens. Two-way ANOVA revealed main effects of cocaine and cocaine x spike on *Grin2b* expression in the nucleus accumbens. (D) Cocaine alone upregulated *Grin2b* mRNA in the prefrontal cortex. Two-way ANOVA revealed a main effect of cocaine on *Grin2b* expression in the prefrontal cortex. Gene of interest relative to *Gapdh* or *Hprt1* mRNA levels in the nucleus accumbens and prefrontal cortex, respectively, were calculated and expressed as relative fold change from the saline-saline control group. Data are expressed as means ± SEM fold change. N = 8-10/group. Significance is denoted as p < 0.05, *; p < 0.01, **.

### Olfaction

Anosmia is a common symptom of both acute COVID-19 and Long-COVID (Kim et al. 2024). Olfaction was tested by quantifying time spent sniffing a neutral odor stimulus (water) or an odor attractant (female rat urine) (Figure 4A,B). Two-way ANOVA revealed main effects of urine (F(3,52) = 3.082, p = 0.0353) and treatment (F(1,52) = 14.41, p = 0.0003), but not interaction between urine x treatment (F(3, 52) = 0.7843, p = 0.5081). Sidak’s multiple comparisons post-hoc analysis of time spent sniffing the odor attractant revealed significantly less time spent sniffing urine for spike-saline versus saline-saline rats (p = 0.0280). These results suggest that olfaction is impaired 2 months after exposure to spike protein.

**Figure 4:**
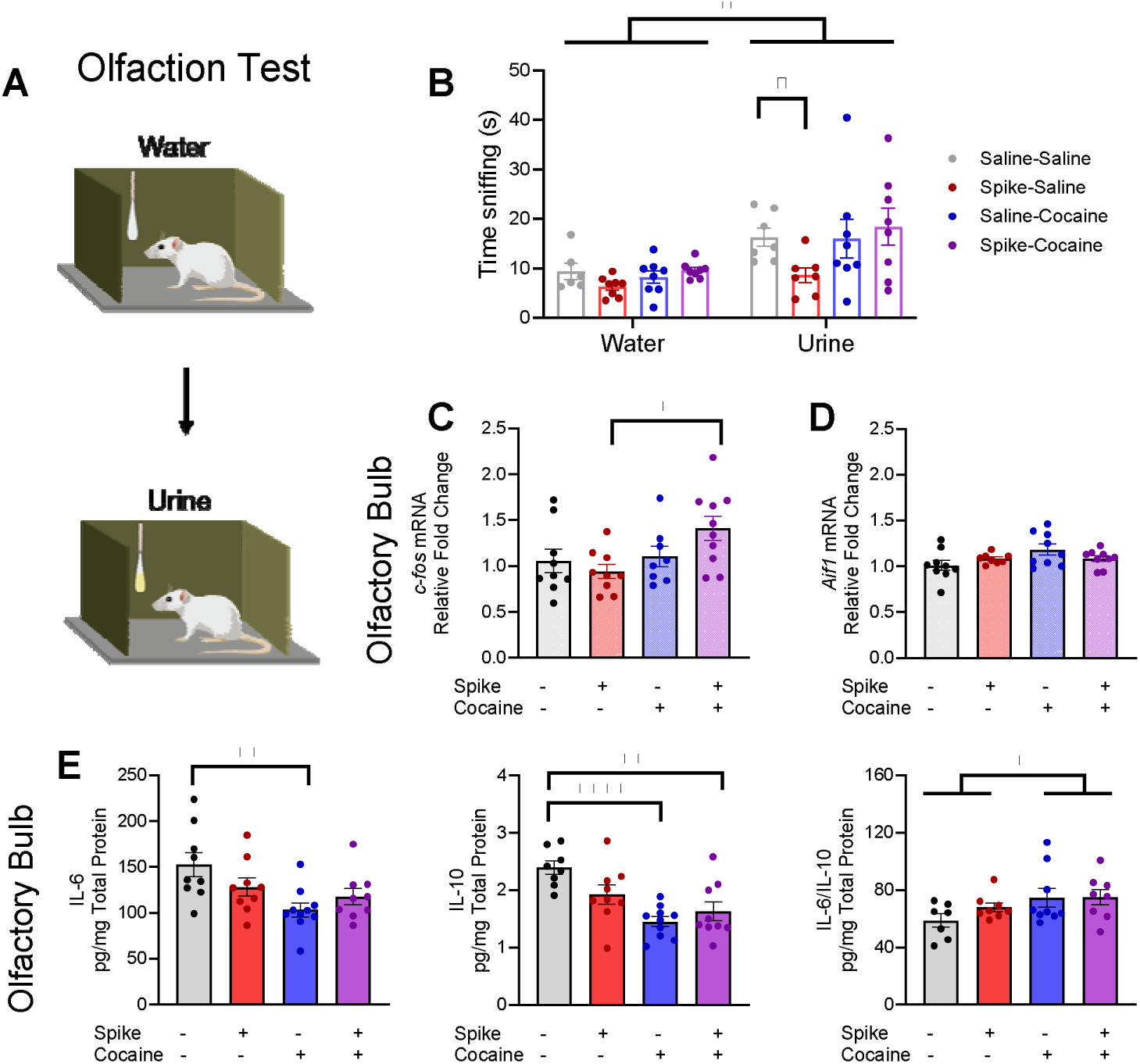
The effect of spike and cocaine on olfaction and gene expression and cytokines in the olfactory bulb. (A) Rats were allowed to freely smell a neutral odor (water) for 3 minutes. After a 45-minute break, animals were placed in a new chamber and allowed to freely smell an odor attractant (female rat urine) for 3 minutes. (B) Time spent sniffing water and urine are shown. Rats injected with spike protein spent less time sniffing an odor attractant compared to saline-saline control animals. Two-way ANOVA revealed main effects of urine and treatment. (C) Relative gene expression for *c-fos* was higher in the spike+cocaine group compared to spike alone in the olfactory bulb. Two-way ANOVA revealed a main effect of cocaine and a trend for an interaction between spike x cocaine. (D) *Aif1* mRNA was unaltered by spike or cocaine in the olfactory bulb. (E) Cocaine alone reduced levels of IL-6 in the olfactory bulb; two-way ANOVA revealed a main effect of cocaine and a trend for an interaction between spike x cocaine on IL-6 levels in the olfactory bulb. IL-10 was downregulated by spike and cocaine; two-way ANOVA revealed main effects of cocaine and interaction between spike x cocaine on IL-10 levels in the olfactory bulb. The ratio of IL-6/IL-10 was higher in rats injected with cocaine, suggesting a pro-inflammatory state; two-way ANOVA revealed a main effect of cocaine on the ratio of IL-6/IL-10 levels in the olfactory bulb. Target genes (C&D) were measured relative to *Gapdh* mRNA, and expressed as fold-change compared with the saline-saline group. Data are expressed as means ± SEM. N = 6-10/group. Significance is denoted as p < 0.05, *; p < 0.01, **; p < 0.0001, ****.

The olfactory bulb relays neuronal signals encoding olfactory information from the nasal olfactory epithelium to the brain’s processing centers for smell (eg, piriform cortex and anterior olfactory nucleus). Animals with a functional olfactory system will exhibit tonic neuronal activity in the olfactory bulb in their home cage, whereas animals with an impacted olfactory system will not (Bepari et al. 2012). Therefore, we quantified *c-fos* expression in the olfactory bulb as a measure of neuronal activity (Figure 4C); tissue was obtained from rats that had been maintained in the home cage without an additional olfactory challenge. Two-way ANOVA of c-fos mRNA revealed a significant effect of cocaine (F(1,32) = 4.524, p = 0.0412), a trend for an interaction between spike x cocaine (F(1,32) = 3.867, p = 0.0580), and no significant effect of spike (F(1,32) = 0.4737, p = 0.4962) on relative c-fos expression in the olfactory bulb. Post-hoc analysis using Sidak’s multiple comparisons test within cocaine groups showed *c-fos* mRNA was higher in the olfactory bulbs of rats exposed to spike-cocaine versus spike alone (p = 0.0109). Since glial cells can express c-Fos under inflammatory conditions (Cruz-Mendoza et al. 2022), the expression of the microglial marker *Aif1* (Iba1) in the olfactory bulb was measured to determine whether the upregulation of *c-fos* could be attributed to microglial activation (Figure 4D). No significant effect of spike was found (F(1,31) = 0.08988, p = 0.7663), however, trends towards significance for cocaine (F(1,31) = 3.41, p = 0.0735) and interaction between spike and cocaine (F(1,31) = 3.312, p = 0.0785) on *Aif1* mRNA expression were observed.

We further investigated the olfactory bulb for potential changes in inflammatory cytokines and chemokines induced by spike and/or cocaine (Figure 4E). For IL-6, two-way ANOVA revealed a significant effect of cocaine (F(1,33) = 8.941, p = 0.0052), a trend for an interaction between spike x cocaine (F(1,33) = 3.791, p = 0.0601), and no significant effect of spike alone (F(1,33) = 0.2532, p = 0.6182). Post-hoc analysis using Sidak’s multiple comparison test revealed significantly lower IL-6 levels in the saline-cocaine versus saline-saline rats (p = 0.0025). For IL-10, two-way ANOVA revealed a significant effect of cocaine (F(1,32) = 20.33, p < 0.0001) and interaction between spike x cocaine (F(1,32) = 5.614, p = 0.0240), and no significant effect of spike alone (F(1,32) = 1.096, p = 0.3031). Post-hoc analysis using Sidak’s multiple comparison test revealed significantly lower IL-10 levels in the saline-cocaine (p < 0.0002) and spike-cocaine (p = 0.0017) versus saline-saline rats. The ratio of IL-6 to IL-10 in the olfactory bulb was elevated by cocaine as revealed by two-way ANOVA; cocaine (F(1,28) = 4.557, p = 0.0417), spike (F(1,28) = 0.8193, p = 0.3731), interaction between spike x cocaine (F(1,28) = 0.7062, p = 0.4078). These results suggest that cocaine drives dynamic changes in cytokines in the olfactory bulb.

### Memory and Anxiety-Like Behaviors

Rats were tested for memory function 8 weeks post spike protein exposure using the novel object recognition task (4-6 days after the last saline or cocaine injection) and the Y-maze (2 days after last saline or cocaine injection). The discrimination index was calculated to determine the rats’ ability to discriminate between the familiar vs. novel objects (Figure 5B). Two-way ANOVA revealed no significant effects of spike (F(1,33) = 0.3311, p = 0.5689), cocaine (F(1,33) = 0.2996, p = 0.5878), or interaction between spike x cocaine (F(1,33) = 0.1317, p = 0.7190) on the discrimination index. We further examined differences in behavior by comparing the total number of interactions with objects during the familiar (Figure 5C) or novel testing phases (Figure 5D). Rats with a history of cocaine exposure had more object interactions during the familiar phase (familiar object 1 + familiar object 2) as revealed by two-way ANOVA; cocaine (Fig 5C; F(1,33) = 5.354, p = 0.0270), spike (F(1, 33) = 0.01541, p = 0.9020), interaction between spike x cocaine (1,33) = 0.1479, p = 0.7030). Total object interactions during the novel phase (familiar object + novel object) also were higher in the cocaine treatment groups as revealed by two-way ANOVA; cocaine (Fig 5D; F(1,33) = 4.752, p = 0.0365), spike (F(1,33) = 0.7258, p = 0.4004), interaction between cocaine x spike (F(1,33) = 0.2509, p = 0.6197).

**Figure 5:**
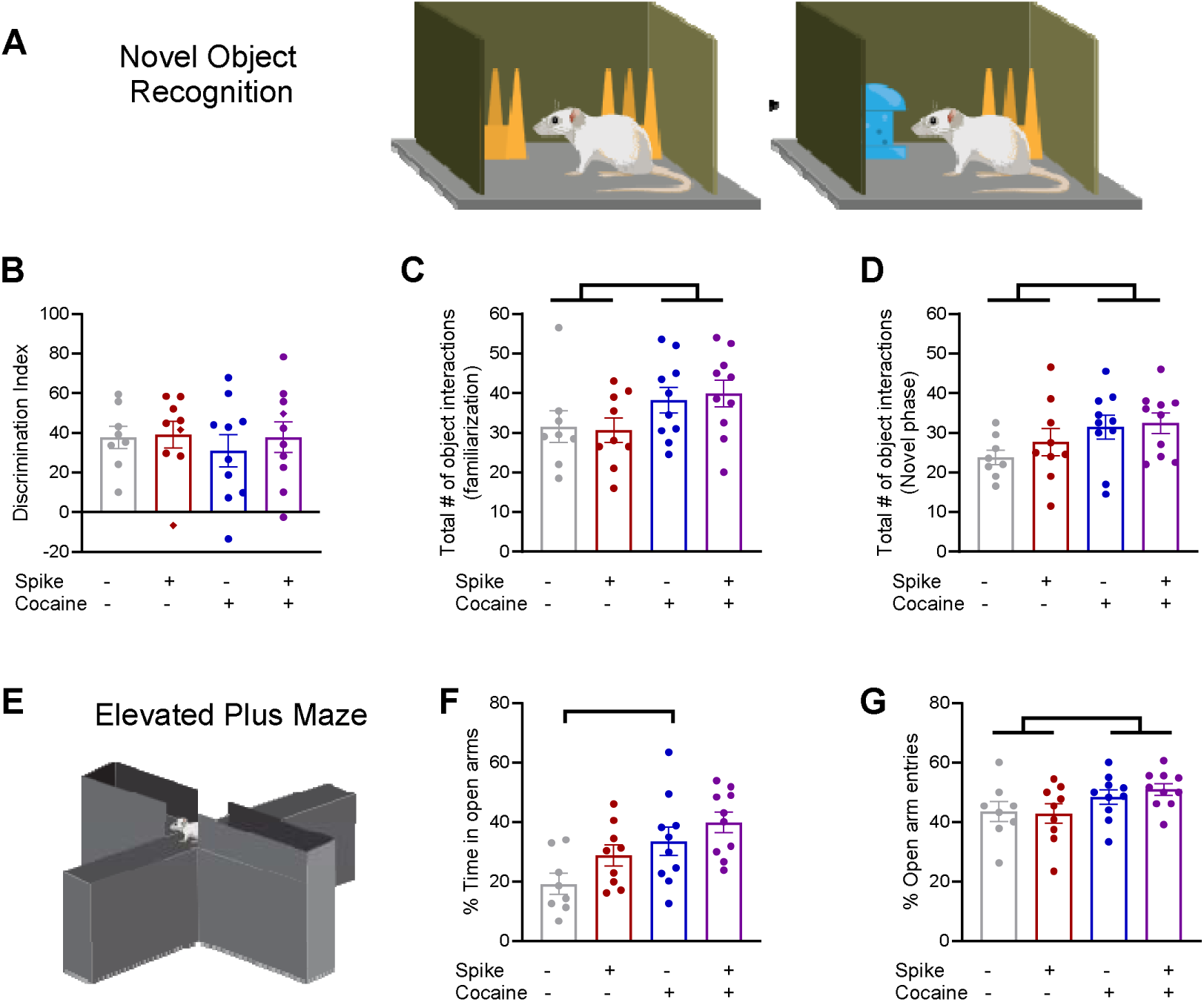
(A) Effect of spike and cocaine on object memory using the novel object recognition task and on anxiety-like behaviors using the elevated plus maze. (B) The discrimination index ([(Time exploring novel – Time exploring familiar)/ (Time exploring novel + Time exploring familiar)] x 100) was used to quantify the ability of the rat to discriminate between the novel and familiar objects. The discrimination index was unaltered by spike and/or cocaine. (C) Cocaine increased the total number of object interactions during the familiarization phase. (D) Cocaine increased the total number of object interactions (novel + familiar) during the novel phase of the task. Two-way ANOVA revealed a main effect of cocaine on both the total object interactions during the familiarization phase as well as during the novel phase of testing. (E) The elevated plus maze was used to assess anxiety-like behavior. Cocaine increased the % time spent in the open arms of the elevated plus maze and (F) the % of open arm entries, suggesting less anxiety-like behaviors in the cocaine rats. Two-way ANOVAs revealed a main effect of cocaine and a trend for an effect of spike on % time in the open arms during the EPM test, and a main effect of cocaine on the % open arm entries during the EPM test. Data are expressed as means ± SEM. N = 8-10/group. Significance is denoted as p < 0.05, *.

The effects of cocaine exposure and/or spike on working spatial memory were tested using the Y-maze (Figure S2). The % spontaneous alternations were higher in the cocaine-treated rats than controls as revealed by two-way ANOVA (F(1,34) = 7.573, p = 0.0094), while neither spike nor cocaine altered the total number of arm entries. The increases in object interactions and spontaneous alternations seen in the cocaine group were unlikely due to differences in locomotor activity, as locomotor activity (total distance traveled, cm, mean velocity, cm/s) did not differ between experimental groups (Figure S3A and S3B) when measured during the habituation phase of the novel object recognition task which occurred 3 or 5 days after the last cocaine or saline injection.

The elevated plus maze (EPM) was used to determine the effects of spike and cocaine on anxiety-like behavior (Figures 5E-G). Two-way ANOVA of the percent time spent in the open arms of the EPM revealed a significant effect of cocaine (F(1,33) = 10.33, p = 0.0029), a trend for a significant effect of spike (F(1,33) = 4.081, p = 0.0516), and no significant interaction between spike x cocaine (F(1,33) = 0.1656, p = 0.6867). Sidak’s multiple comparisons test revealed that the saline-cocaine group spent significantly more time in the open arms than the saline-saline group (p = 0.0332) (Fig 5F). Furthermore, the percent open arm entries was higher in the cocaine groups as revealed by two-way ANOVA; cocaine (F(1,33) = 5.510, p = 0.0251), spike (F(1,33) = 0.1207, p = 0.7305), interaction between spike x cocaine (F(1,33) = 0.3451, p = 0.5609)(Fig 5G).

### Spike or Cocaine may produce an “impulsive-like” phenotype

The elevated plus maze is commonly used to assess whether an animal exhibits anxiety-like behavior based on the time spent and entries into the open arms of the maze (Lapiz-Bluhm et al. 2008). However, an increase in time spent exploring the open arms of an EPM as well as latency to explore a new environment has been interpreted to indicate an impulsive-like behavioral phenotype (Rico et al. 2017). To further explore the possibility that spike and/or cocaine may produce an “impulsive-like” phenotype, the latency to the first and second open arm entries during the EPM task was measured (Figure 6A and B). Spike reduced the latency to the first open arm entry as revealed by two-way ANOVA; spike (F(1,27) = 10.83, p = 0.0028), cocaine (F(1,27) = 2.751, p = 0.1088), interaction between spike x cocaine (F(1,27) = 2.282, p = 0.1425). Sidak’s multiple comparisons test revealed significantly shorter time to first arm entry in the spike-saline vs. saline-saline groups (p = 0.0078)(Fig 6A). Cocaine reduced the latency to the second open arm entry as revealed by two-way ANOVA; cocaine (F(1,32) = 6.608, p = 0.0150), spike (F(1,32), p = 0.1909, p = 0.6651), interaction between spike x cocaine (F(1,32) = 0.1662, p = 0.6863)(Fig 6B). Although this is not a rigorous measurement of impulsivity, the results suggest that rats exposed to cocaine or spike protein exhibited more impulsive-like behaviors than untreated controls.

**Figure 6:**
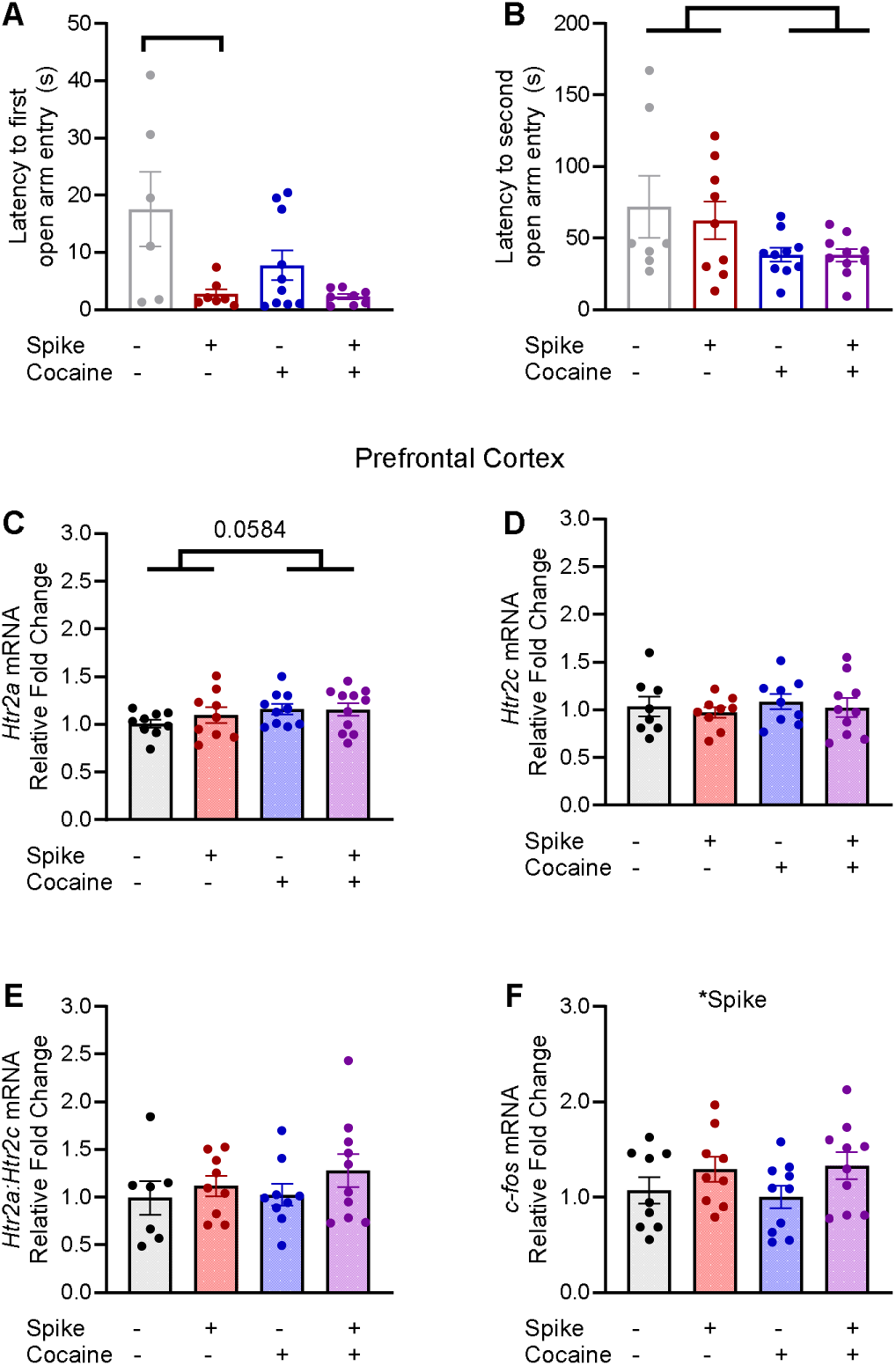
Spike or cocaine may produce an “impulsive-like” phenotype. (A) Latencies to enter an open arm of the elevated plus maze were recorded. Rats injected with spike protein entered an open arm more quickly than saline-saline controls. Two-way ANOVA revealed a main effect of spike on the latency to the first open arm entry during the EPM test. (B) Cocaine significantly reduced the latency to the second entry into an open arm of the plus maze. Two-way ANOVA revealed a main effect of cocaine on the latency to the second open arm entry during the EPM test. (C) *Htr2a* mRNA in the prefrontal cortex was non-significantly elevated in cocaine-treated groups. Two-way ANOVA revealed a trend for a main effect of cocaine on *Htr2a* expression in the prefrontal cortex. (D) *HTR2c* expression was not different between groups. (E) Neither cocaine nor spike altered the ratio of *Htr2a/Htr2c* mRNA in the prefrontal cortex. (F) *c-fos* mRNA in the prefrontal cortex was higher in rats injected with spike protein; two-way ANOVA revealed a main effect of spike on *c-fos* expression. Gene of interest relative to *Hprt1* mRNA of each sample was measured and converted to fold change relative to the saline-saline controls. Data are expressed as means ± SEM. N = 6-10/group. Significance is denoted as p < 0.05, *; p < 0.01, **.

The role of the serotonin receptors 5-HT2aR and 5-HT2cR in the prefrontal cortex in mediating both impulsive and cocaine-induced impulsive behaviors is well evidenced (Cunningham et al. 2013; Anastasio et al. 2015). Therefore, *Htr2a* (5-HT2a) and *Htr2c* (5-HT2c) mRNAs in the prefrontal cortex were quantified. There was a trend for cocaine to upregulate *Htr2a* expression as revealed by two-way ANOVA (Fig 6C); cocaine (F(1,34) = 3.836, p = 0.0584), spike (F(1,34), = 0.9844, p = 0.3281), interaction between spike x cocaine (F(1,34) = 0.1944, p = 0.6621). Two-way ANOVA revealed no significant effect of spike or cocaine on *Ht2rc* (Fig 6D); cocaine (F(1,32) = 0.3347, p = 0.5670), spike (F(1,32) = 0.4977, p = 0.4856), interaction between spike x cocaine (F(1,32) = 0.0001604, p = 0.9900). The balance of 5-HT2aR to 5-HT2cR levels is important in modulating impulsive behavior, such that higher 5HT-2aR:5-HT2cR ratios are associated with higher impulsivity (Anastasio et al. 2015). Here, neither spike nor cocaine altered the ratio of *Htr2a/Htr2c* in the prefrontal cortex as revealed by two-way ANOVA (Figure 6E); cocaine (F(1,31) = 0.4352, p = 0.5143), spike (F(1,31) = 1.644, p = 0.2093), interaction between spike x cocaine (F(1,31) = 0.1940, p = 0.6627). In addition to 5-HT2 receptor balance, neuronal activity in the prefrontal cortex is hypothesized to play a role in impulsivity (Anastasio et al. 2015). Therefore, we measured *c-fos* expression in the prefrontal cortex (Figure 6F). *c-fos* mRNA was higher in the prefrontal cortex of spike-injected rats as revealed by two-way ANOVA; spike (F(1,34) = 4.302, p = 0.0457), cocaine (F(1,34) = 0.1575, p = 0.9009), interaction between spike x cocaine (F(1,34), = 0.1709, p = 0.6819).

## Discussion

The present study explored the long-term effects of SARS-CoV-2 spike protein in the setting of chronic cocaine exposure on Long-COVID-like outcomes in a rat model. A single intravenous injection of spike protein (500 ng/kg) was administered to model an acute infection of SARS-CoV-2 virus. The peak level of spike antigen during the infectious phase of SARS-CoV-2 viral infection is not currently known. Based on the molecular weight of spike protein (homotrimer), the average number of spike proteins per virion, and the viral load in the lungs of macaques (Sender et al. 2021; Rebelo et al. 2022), we estimated a peak spike concentration of 40.35 – 4,034 ng/kg. Based on this estimation, a dose of 500 ng/kg spike was chosen for this study. Importantly, the Wuhan strain of spike has a nanomolar affinity (Kd = 2.76 nM) for the rat ACE2 receptor (Yao et al. 2021). While the Kd is higher for rACE2 compared to the human ACE2 receptor (Kd = 0.71 nM), Sprague Dawley rats were found to be susceptible to infection by the Wuhan strain of the SARS-CoV-2 virus (Yu et al. 2022). Future work using rats as a model to study the effects of spike may require higher doses of spike to account for the lower affinity of spike for rACE2 compared to hACE2. For the field of Long-COVID research to progress and provide translational findings, there is a critical need to quantify the spike protein load produced by the SARS-CoV-2 virus to establish a clinically relevant model.

The first major finding in this study was the persistent impact of spike and cocaine on body weight gain. Chronic cocaine use in humans reduces body fat and weight (Ersche et al., 2013), while separately, acute COVID-19 (regardless of Long-COVID status) leads to lean body mass (Ersche et al. 2013; Atieh et al. 2024). A preclinical study found that mice (K18-hACE2) fed an alcohol diet and exposed to the S1 subunit of the spike protein had lower body weights compared mice exposed to spike (S1) or alcohol alone (Solopov et al. 2022). The present study provides the first evidence that exposure to the spike protein in the setting of chronic cocaine exposure produced a persistent reduction in weight gain.

SARS-CoV-2 and cocaine independently increase inflammatory signatures, including the microgliosis activation marker Iba1 and the astroglia activation marker GFAP (Scofield et al. 2016; Linker et al. 2019; Serrano et al. 2022; Soung et al. 2022; Bark et al. 2023; Rong et al. 2024). Elevated Iba1 protein levels were found in postmortem samples of the medulla of humans with acute COVID-19, the hippocampus and medulla of hamsters intranasally infected with SARS-CoV-2 virus (Soung et al. 2022), and the cortex of mice injected with the spike protein into the skull marrow (Rong et al. 2024). Likewise, induction of GFAP is associated with SARS-CoV-2. For example, elevated blood levels of GFAP are associated with mild cognitive dysfunction in persons with Long-COVID (Bark et al. 2023), and cerebral spinal fluid (CSF) levels of GFAP are elevated in Long-COVID, but not acute COVID-19, patients (Rong et al. 2024). Regarding cocaine, Iba1 and GFAP protein levels are elevated in various brain regions (nucleus accumbens, hippocampus, dentate gyrus) of mice injected with cocaine (Fattore et al. 2002; Liao et al. 2016; Lewitus et al. 2016; Linker et al. 2019). Here, we found divergent effects of cocaine on the regulation of the gene encoding Iba1, *Aif1*, with *Aif1* mRNA expression downregulated in the hippocampus and a trend for upregulation in the prefrontal cortex (Figure 1C and D). Prior work suggests that *Aif1* expression does not necessarily correlate with Iba1 protein levels. A previous report wherein the proinflammatory molecule lipopolysaccharide was injected into mice found that while Aif1 gene expression was downregulated in the hippocampus, immunohistochemistry staining showed elevated Iba1 protein (Silverman et al. 2015). Our results suggest that spike and cocaine may affect microglial activation; future work is necessary to quantify Iba1 protein levels and microglial morphology in these brain regions.

The second major finding of this study was that the combination of spike and cocaine (but neither alone) increased mechanical sensitivity which was correlated with IL-10 levels. Heightened paw sensitivity, as shown by lower withdrawal thresholds in the Von Frey test, was associated with lower IL-10 levels in the prefrontal cortex and hippocampus than controls (Figure 2B, E, F). Chronic musculoskeletal pain is reported by people with Long-COVID and is linked to dysregulation of inflammatory cytokine levels (Khoja et al. 2024). For example, pain severity in persons with Long-COVID is negatively correlated with serum IL-10 levels (Bussmann et al. 2022). Evidence for musculoskeletal pain or mechanical allodynia due to cocaine use is rare (de Souza et al. 2017), although cytokine regulation is not. Lower IL-10 and higher IL-6 levels in serum of cocaine users compared to controls have been demonstrated (Moreira et al. 2016). In addition to the negative correlation observed between mechanical sensitivity and IL-10 levels in the present study, we observed a shift towards a pro-inflammatory state, as evidenced by an increase in the ratio of IL-6/IL-10 (You et al. 2011; Gholamnezhad et al. 2014; Xie et al. 2015), in the hippocampus of rats that received spike and cocaine compared to the untreated control group. Taken together, these results support that co-exposure to cocaine and spike produced late-onset mechanical allodynia associated with dysregulation of IL-10 in levels the brain. A future goal will be to determine the therapeutic potential for IL-10 in alleviating musculoskeletal pain due to Long-COVID with or without co-occurring cocaine use. This direction is supported by the finding that intrathecal administration of IL-10 alleviates mechanical allodynia in multiple rodent models of pain (Laumet et al. 2020; Huang et al. 2022).

We further explored possible molecular disturbances that may underlie cocaine- and spike-induced mechanical allodynia by measuring changes in gene expression for the GluA1 (*Gria1)* subunit of the glutamatergic AMPA receptor and the GluN2B (*Grin2b)* subunit of the NMDA receptor in the prefrontal cortex and nucleus accumbens. A study in rats found that persistent tactile allodynia is associated with abnormal brain activity in the prefrontal areas and nucleus accumbens, suggesting these areas may encode the lasting effect of tactile allodynia following neuropathic injury (Chang et al. 2017). In the present study, *Gria1* expression was upregulated in the prefrontal cortex by spike or cocaine. This finding agrees with prior work showing elevated GluA1 levels in the prefrontal cortex of mice with inflammatory pain induced by complete Freud’s adjuvant (Li et al. 2017). Other studies in rodent inflammatory pain models accompanied by mechanical allodynia have observed upregulation of the glutamatergic NMDA receptor subunit GluN2B in the anterior cingulate cortex (Li et al. 2017; Fan et al. 2018). Upregulation of GluN2B containing NMDA receptors in the anterior cingulate cortex of mice was observed in a model of neuropathic pain using chronic constrictive injury, while knockdown of GluN2B in anterior cingulate cortex neurons prevented chronic constrictive injury-induced mechanical allodynia (Wang et al. 2021). We observed a downregulation of *Grin2b* mRNA expression in the nucleus accumbens and an upregulation in the prefrontal cortex of rats exposed to cocaine with or without spike (Figure 3C and D). Further investigation of AMPA and NMDA receptor subunits in the prefrontal cortex and nucleus accumbens may provide key insights into the pathology underlying mechanical allodynia produced by spike protein in the presence of cocaine.

The third major finding of this study was that spike alone reduced time spent sniffing an odor attractant months after spike exposure, recapitulating a prominent feature of both acute COVID-19 and Long-COVID (Kim et al. 2024). The cause of SARS-CoV-2-induced anosmia is unknown, however, increases in immune cells in the olfactory epithelium and downregulation of odorant receptor genes in olfactory sensory neurons are observed in multiple animal models infected with SARS-CoV-2 (Zheng et al. 2021; Tsukahara et al. 2023). Regarding the spike protein alone, anosmia is observed 3 days after intranasal exposure to spike protein in zebrafish (Kraus et al. 2022). Intriguingly as related to our study, a retrospective study of people with Long-COVID using the TriNetX database identified a higher risk for sensory impairment in persons with co-occurring substance use disorders compared to those without a substance use disorder (Tsai et al. 2024). In the present study, only the rats injected with spike alone had impaired olfaction. We further explored the potential impact of spike and cocaine on olfactory neurons by determining basal neuronal activity in the olfactory bulb by measuring *c-fos* expression. Basal activation of olfactory sensory and relay neurons in the olfactory bulb occurs when an animal is in its home-cage (Bepari et al. 2012), and *c-fos* expression in the olfactory bulb is an indicator of intact olfaction processes (Weiler et al. 2006; Bepari et al. 2012). In contrast with our hypothesis, we observed higher *c-fos* mRNA in rats treated with chronic cocaine and spike (Figure 4C). To investigate whether the increase in *c-fos* could be related to activation of glial cells which also express *c-fos* (Cruz-Mendoza et al. 2022), we measured the expression of the microglial marker *Aif1* in the olfactory bulb (Figure 4D). Neither spike nor cocaine altered *Aif1* expression in the olfactory bulb. This study presents the first evidence that mammalian olfaction is impaired months after exposure to spike protein. Furthermore, this study indicates that impaired olfaction can occur after a non-intranasal route of administration of spike, in this case intravenous injection.

Previous work reported that intracerebroventricular injection of spike protein (6.5 μg/mouse) produces deficits in object and spatial memory 30 and 45 days following spike exposure (Fontes-Dantas et al. 2023). Here, spike (500 ng/kg, i.v.) had no effect on object or spatial memory when tested 60 days after injection (Figure 5A-D, Figure S2). The discrepancy between the outcomes of the two studies is likely due to the dose of spike used (6.5 ug/mouse vs 500 ng/kg rat body weight) considering previous dose dependent studies (Fontes-Dantas et al. 2023). In addition to memory deficits, anxiety is a common symptom of Long-COVID (Nalbandian et al. 2023). Intra-skull marrow injection of the S1 subunit of the spike protein (N501Y) produces an anxiety-like phenotype 3 days, but not 28 days, following injection, as evidenced by reduced time spent in the center of an open field (Rong et al. 2024). No such anxiogenic effect was found in the present study using the open field test when measured 60 days post spike injection in rats. In fact, in the elevated plus maze (EPM) test, cocaine-exposed animals spent more time in the open arms, with a similar non-significant trend for the spike group (Figure 5E).

Our interpretation of the EPM results leads us to the final major finding of this study, which is that spike and cocaine may produce an “impulsive-like” phenotype as evidenced by reduced latencies to the first and second open arm entries during the EPM test. While the EPM is intended to assess anxiety-like phenotypes, some behaviors on the EPM have been found to correlate with impulsivity (Rico et al. 2017). For example, rats with a preestablished high impulsivity phenotype have reduced latency to first open arm entry compared to rats with a low impulsivity phenotype (Molander et al. 2011). Here, spike reduced the latency to the first open arm entry of the EPM, while cocaine reduced the latency to the second open arm entry (Figure 6 A-B). It is not surprising that long-term cocaine administration would produce a possible “impulsive-like” phenotype, as cocaine-driven impulsivity is well-established in both humans and rodent models (Verdejo-García et al. 2007; Simon et al. 2007; Mendez et al. 2010). Cocaine-driven impulsivity is causally linked to upregulation of 5-HT2a receptors relative to 5-HT2c receptors in the rat prefrontal cortex (Cunningham et al. 2013; Anastasio et al. 2015). We found a trend for cocaine-induced upregulation of *Htr2a* expression (Figure 6C). We also found that *c-fos* expression was higher in the prefrontal cortex of rats injected with spike protein two months earlier (Figure 6F), possibly indicating heightened neuronal activity. A recent report using electrophysiology found that bath application of SARS-CoV-2 spike protein produced hyperexcitable changes in pyramidal neurons in layers V-VI of medial prefrontal cortex in rats, which was exacerbated by a history of cocaine self-administration (Cassoday 2025). Activation of 5-HT2a receptors on layer V pyramidal neurons, which increases neuronal excitability and the frequency of spontaneous excitatory postsynaptic currents, is hypothesized as the mechanism by which cocaine-induced upregulation of 5-HT2a to 5-HT2c receptors in the prefrontal cortex produces impulsive behavior (Anastasio et al. 2015; Berthoux et al. 2019). The effect of spike and cocaine on neuronal excitability in the prefrontal cortex presents an interesting avenue for future research that may help to elucidate mechanisms of altered impulsive-like behaviors. In addition, our future work will address whether spike and/or cocaine induces an impulsive phenotype using a direct measure of impulsivity such as the delay discounting task (Mar and Robbins 2007).

The present study sought to explore the effects of SARS-CoV-2 spike protein in the setting of chronic cocaine exposure on Long-COVID-like outcomes in the rat. There are limitations to this study. First, the dose of spike used was based on an estimation of spike load produced during viral infection. Future work is needed to quantify spike protein during infection to ensure a clinically relevant dose of spike is used in preclinical studies. Second, the effects of the spike protein alone and not SARS-CoV-2 virus as a whole were tested. Although detection of viral RNA is rare in Long-COVID after initial infection has cleared (Salman et al. 2021), it does not eliminate the possibility that other factors associated with viral infection may contribute to Long-COVID. Lastly, the present study was performed using only young adult male rats. Long-COVID symptoms and brain alterations present with sex-specific patterns and are worsened with age (Reas et al. 2025). Future work is needed to determine how cocaine impacts spike-induced Long-COVID outcomes in the context of sex and age.

In summary, cocaine negatively impacts several Long-COVID-like outcomes in a spike-injected rat model of Long-COVID. Specifically, spike plus cocaine reduced weight gain, induced mechanical allodynia, and promoted a proinflammatory state in the hippocampus. Further, olfaction was impaired months after exposure to spike, recapitulating a prominent feature of Long-COVID. Finally, reduced latency to enter open arms in the elevated plus maze following spike or cocaine exposure suggests an “impulsive-like” phenotype in these animals. Our findings highlight the need for further research into the co-occurring effects of SARS-CoV-2 and substance use disorders and increased clinical observations of people with co-occurring cocaine use disorder and Long-COVID. Lastly, these findings support the use of a spike injection as a model of Long-COVID in the rat.

## Acknowledgments

We thank Kofi Osei-Abrefa Ayensu for his assistance with the study. This work was funded by NIH grants R01DA054921, P30DA013429, T32DA007237 to Ellen Unterwald, T34GM136494 to Spiridoula Matsika (Daniel Lopez), and R25NS130632 to Lisa Briand (Flourish Anwuri).

## SUPPLEMENTARY DATA

As detailed in the Methods Section, rats received intermittent chronic cocaine administration and a single intravenous injection of the SARS-CoV-2 spike protein (500 ng/kg). Two months following spike administration, Long-COVID-like outcomes were assessed.

### Results

Supplemental Figure S1 shows IL-6 and IL-10 levels, as well as the ratio of IL-6:IL-10, in the serum of rats two months after exposure to spike. IL-6 and IL-10 levels in the serum were unchanged two months after exposure to spike with or without chronic cocaine administration. The ratio of IL-6:IL-10 was higher in rats exposed to cocaine with or without spike (Fig S1C).

Supplemental Figure S2 shows % spontaneous alternations and total number of arm entries during the Y-maze test performed two months after exposure to spike with or without chronic cocaine. Rats that received cocaine had higher % spontaneous alternations than those without cocaine (Fig S2A), while total number of arm entries was unaffected by spike or cocaine exposure (Fig S2B).

Supplemental Figure 3 shows locomotor activity and performance during the open-field test as a measure of anxiety-like behaviors. Neither spike nor cocaine affected total distance traveled (Fig S3A), mean velocity (Fig S3B), time spent in the center (Fig S3C), or total number of center entries (Fig S3D) during the open field test.

**Figure S1:**
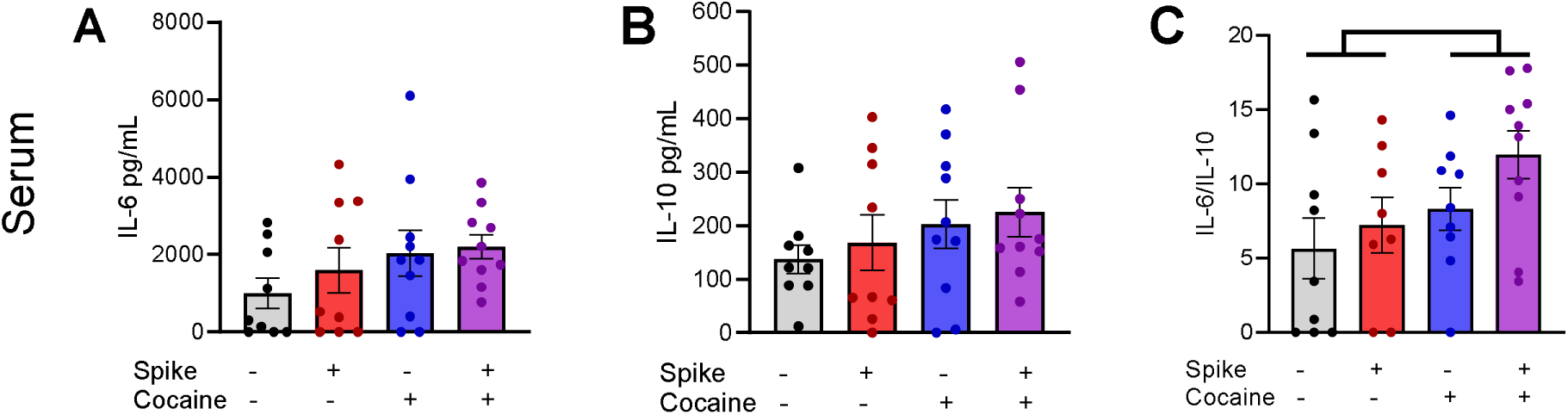
IL-6 and IL-10 were quantified in serum obtained two months post spike protein administration. (A) No differences were observed in IL-6 levels in the serum of rats exposed to spike or cocaine. Two-way ANOVA: spike (F(1,34) = 0.6285, p = 0.4334), cocaine (F(1,34) = 2.860, p = 0.1000), spike x cocaine (F(1,34) = 0.1891, p = 0.6664). (B) No differences were observed in IL-10 levels in the serum following exposure to spike or cocaine. Two-way ANOVA: spike (F(1,34) = 0.3663, p = 0.5490), cocaine (F(1,34) = 1.936, p = 0.1731), spike x cocaine (F(1,34) = 0.0098, p = 0.9218). (C) The ratio of IL:6-IL-10 in the serum was higher in cocaine-exposed groups with or without spike compared with saline controls. Two-way ANOVA: spike (F(1,32) = 0.3463, p = 0.5604), cocaine (F(1,32) = 4.406, p = 0.0438), spike x cocaine (F(1,32) = 2.206, p = 0.1472). Data are expressed as means ± SEM. N = 9-10/group. Significance is denoted as p < 0.05, *.

**Figure S2:**
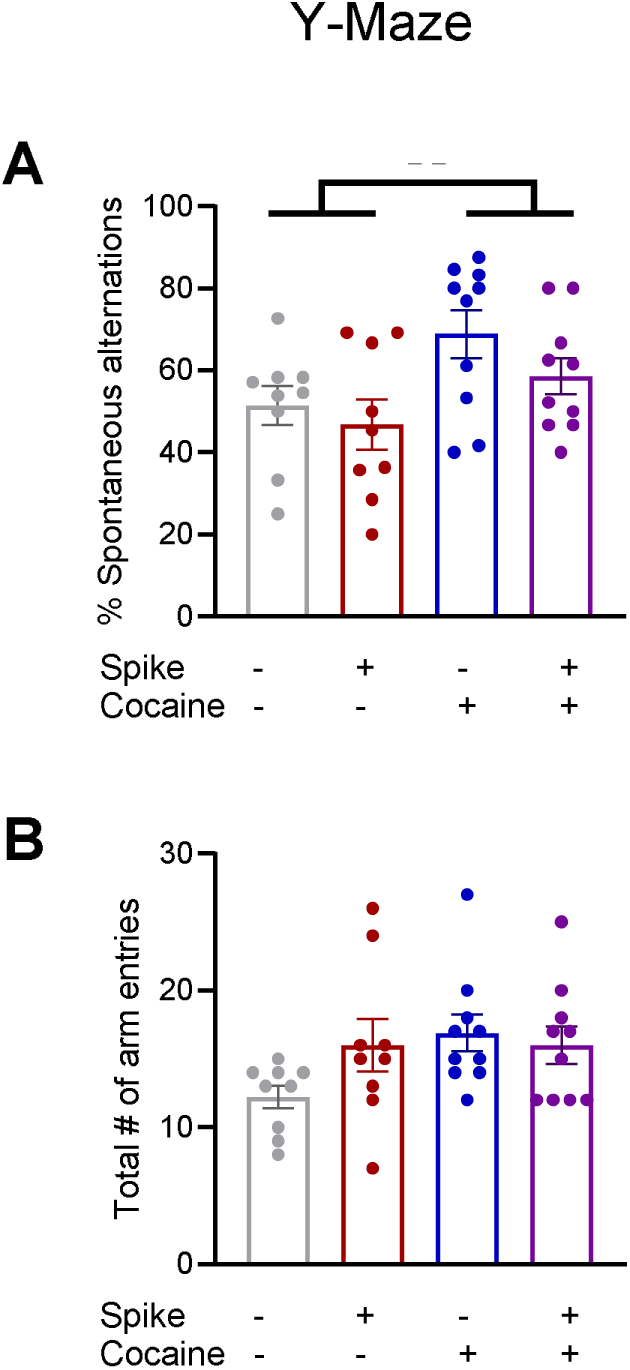
Effects of spike and/or cocaine on spatial working memory were assessed using the Y-maze. (A) The % of spontaneous alternations during the Y-maze task were greater in rats injected with cocaine compared with saline controls; two-way ANOVA: spike (F(1,34) = 1.972, p = 0.1693), cocaine (F(1,34) = 7.573, p = 0.0094), spike x cocaine (1,34) = 0.2743, p = 0.6040). (B) Neither cocaine nor spike altered the total number of arm entries during the Y-maze task. Two-way ANOVA: spike (F(1,34) = 1.022, p = 0.3192), cocaine (F(1,34) = 2.701, p = 0.1095), spike x cocaine (F(1,34) = 2.701) = 0.1095). Data are expressed as means ± SEM. N = 9-10/group. Significance is denoted as p < 0.01, **.

**Figure S3:**
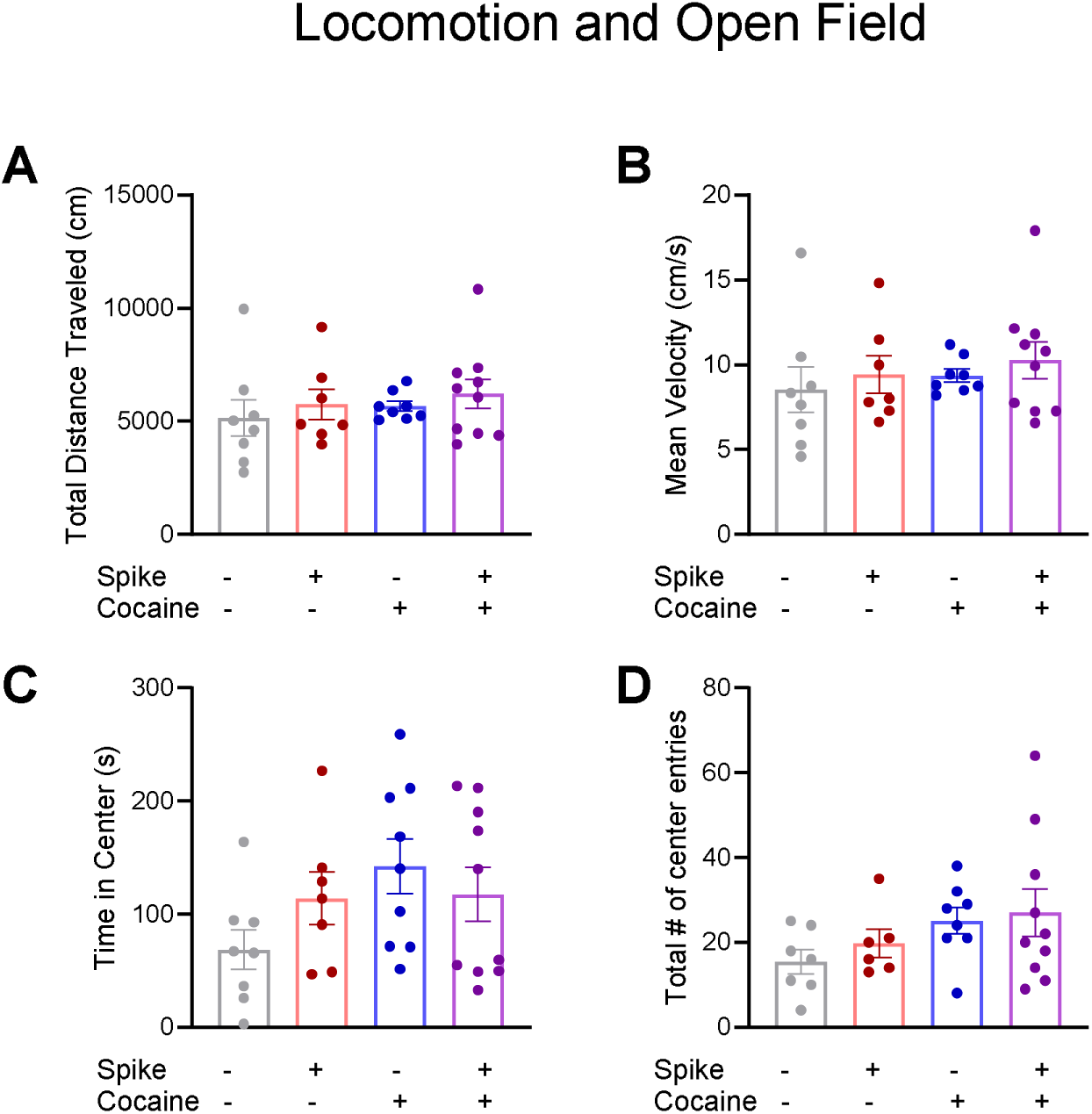
Anxiety-like behavior and locomotor activity were assessed using the open field test. Neither spike or cocaine altered locomotor activity (A. total distance traveled or B. mean velocity) or anxiety-like behavior (C. time spent or D. entries into the center zone). Data are expressed as means ± SEM. N = 9-10/group.

## Notes

### Competing Interest Statement

The authors have declared no competing interest.

